# Productive and latent HIV infections originate in resting CD4^+^ T cells

**DOI:** 10.1101/2022.08.26.505461

**Authors:** Stephen W. Wietgrefe, Jodi Anderson, Lijie Duan, Peter J. Southern, Paul Zuck, Guoxin Wu, Bonnie J. Howell, Cavan Reilly, Eugène Kroon, Suthat Chottanapund, Supranee Buranapraditkun, Carlo Sacdalan, Nicha Tulmethakaan, Donn J. Colby, Nitiya Chomchey, Peeriya Prueksakaew, Suteeraporn Pinyakorn, Rapee Trichavaroj, Denise Hsu, Sandhya Vasan, Sopark Manasnayakorn, Mark de Souza, Sodsai Tovanabutra, Alexandra Schuetz, Merlin L. Robb, Nittaya Phanuphak, Jintanat Ananworanich, Timothy W. Schacker, Ashley T. Haase

## Abstract

Productively and latently HIV-infected cells are the source of virus that respectively establishes and sustains systemic infections and the reservoir in which HIV persists and rebounds when anti-retroviral therapy (ART) is interrupted. While infected activated CD4^+^ T cells are thought to be the principal source of HIV production, and reversion of activated infected cells to a resting state as the major pathway to establishment of the latently infected cell reservoir, we now show that in the earliest stages of detectable HIV infection in the lymphoid tissue reservoir, infection of resting CD4^+^ T cells establishes the first populations of both productively and latently infected cells. We further show that the early infection of resting T cells reflects their predominance in lymphoid tissues and the expression of pTEFb in vivo in resting T cells to support their infection. The immediate establishment of productively and latently infected cell populations enable HIV to propagate and persist, and generates reservoirs from which infection can rebound despite instituting ART at the earliest stage of detectable infection.

## HIV and SIV infections in activated and resting CD4 T cells

From the beginnings of HIV research, the virus has been propagated in vitro mainly in cultures of activated CD4^+^ T cells^1^, leading to the prevailing view that activated CD4^+^ T cells are the major source of virus production in vivo. This conclusion is particularly well supported by studies of the dynamics of the response to anti-retroviral therapy (ART), which blocks new infections to result in first phase decay of activated infected cells in peripheral blood and lymphoid tissues (LT)^2, 3^. The slower second and later stages of decay and persistence of HIV infection are thought to reflect the establishment of a reservoir of transcriptionally silenced^4^, latently infected resting CD4^+^ T cell populations^5-8^ thought to arise when infected activated CD4^+^ T cells, *pari passu* their uninfected counterparts, return to an immunologically resting state^9^.

However, there is compelling in vivo evidence in studies of the first days and weeks of SIV infection following mucosal infection that infected resting CD4^+^ T cells initially predominate both at the portal of entry and in LT to which virus spreads^10-12^, and these infected resting T cells largely account for SIV production at the earliest stages of infection^13^. These studies of SIV infection in the NHP model also revealed correlations that support a model in which target cell availability determines the cell type in which virus replicates^11^. Initially, resting CD4^+^ T cell populations are the predominant susceptible target cell and the predominant cell in which SIV replicates. As resting CD4^+^ T cells are depleted, and immune activation provides populations of activated T cells, these activated populations provide the target cells that increasingly support expanding virus production^12^.

Could the initial events in HIV infection be similar to SIV infection in which the preponderant target cell availability of resting CD4^+^ T cells dictates establishment of productive infections in HIV-infected resting T cells? And, if so, could infected resting T cells, rather than infected activated T cells, be the source of the latently infected cell population? The RV254 study of acute HIV infection that included the earliest detectable stage of infection (Fiebig 1) provided an opportunity to answer this question.

### Thailand cohort of the earliest stages of HIV infection

In the RV254 study, participants with acute HIV are recruited at the Thai Red Cross AIDS Research Centre where blood samples of clients presenting for voluntary counseling and testing were screened by nucleic acid or sequential immunoassay within 1-2 days of sample collection^14^. Participants found to have acute HIV infection (Fiebig stages 1-5) were asked to enroll in a study in which lymph node (LN) biopsies were obtained in a subset of consenting volunteers. For the studies reported here of the earliest detectable stage of HIV infection, we analyzed LN biopsies from nine participants in Fiebig 1 to determine the cell types and target cell activation status in which HIV first replicates. In the studies of the available LN suspensions that we assayed for reactivatable latently infected cells in Fiebig 1, we also assayed LN suspensions from one participant in Fiebig 4 and one participant in Fiebig 5 for comparison (demographic and Fiebig staging information for the 11 participants is given in supplemental Table 1).

### HIV RNA^+^ and virus-producing cells are T cells with a resting phenotype in Fiebig 1

We screened for HIV RNA^+^ and virus-producing cells, using RNAscope in situ hybridization (ISH) and probes specific for the uniformly AE or AE/B recombinant-clade HIV infections in these subjects, to identify HIV RNA^+^ cells. After developing a method for ISH with ELF97 substrate to clearly visualize HIV virions in virus-producing cells^15^, we subsequently screened LN tissue sections for HIV-producing cells, and detected HIV RNA^+^ virus producing cells in 9/9 subjects, either as isolated single cells; two cells together, consistent with cell-to-cell transmission; or two cells in close spatial proximity (Figure 1A). By comparing subjacent sections, we found that 100 percent of HIV RNA^+^ cells were also virus^+^, i.e., visibly producing virus (Table 1) to thereby assign virus production in these LN samples to HIV RNA^+^ virus-producing cells.

**Table 1.** Phenotype of HIV RNA^+^ Virus-Producing Cells in LN in Fiebig 1.

| **Subject ID** | **HIV RNA<sup>+</sup>/Virus<sup>+</sup>**<br>**(percent)** | **HIV RNA<sup>+</sup>/CD3<sup>+</sup>**<br>**(percent)** | **HIV virus<sup>+</sup>/CD4<sup>+</sup>**<br>**(percent)** | **HIV virus<sup>+</sup>/CD25<sup>+</sup>**<br>**(percent)** |
| --- | --- | --- | --- | --- |
| 2265 | 100 | 100 | - | - |
| 2391 | 100 | 100 | 88 | 0 |
| 2397 | 100 | 100 | 100 | - |
| 2246 | 100 | 100 | 89 | 0 |
| 2540 | 100 | 98 | 92 | 0.63 |
| 2451 | 100 | 100 | 100 | 0 |
| 2563 | 100 | 100 | 100 | - |
| 2585 | 100 | 100 | 96 | 0 |
| 2494 | 100 | - | 100 | 0 |

Key: Negative (-) sign indicates that HIV<sup>+</sup> cells were not detected for phenotypic analysis.

**Figure 1.**
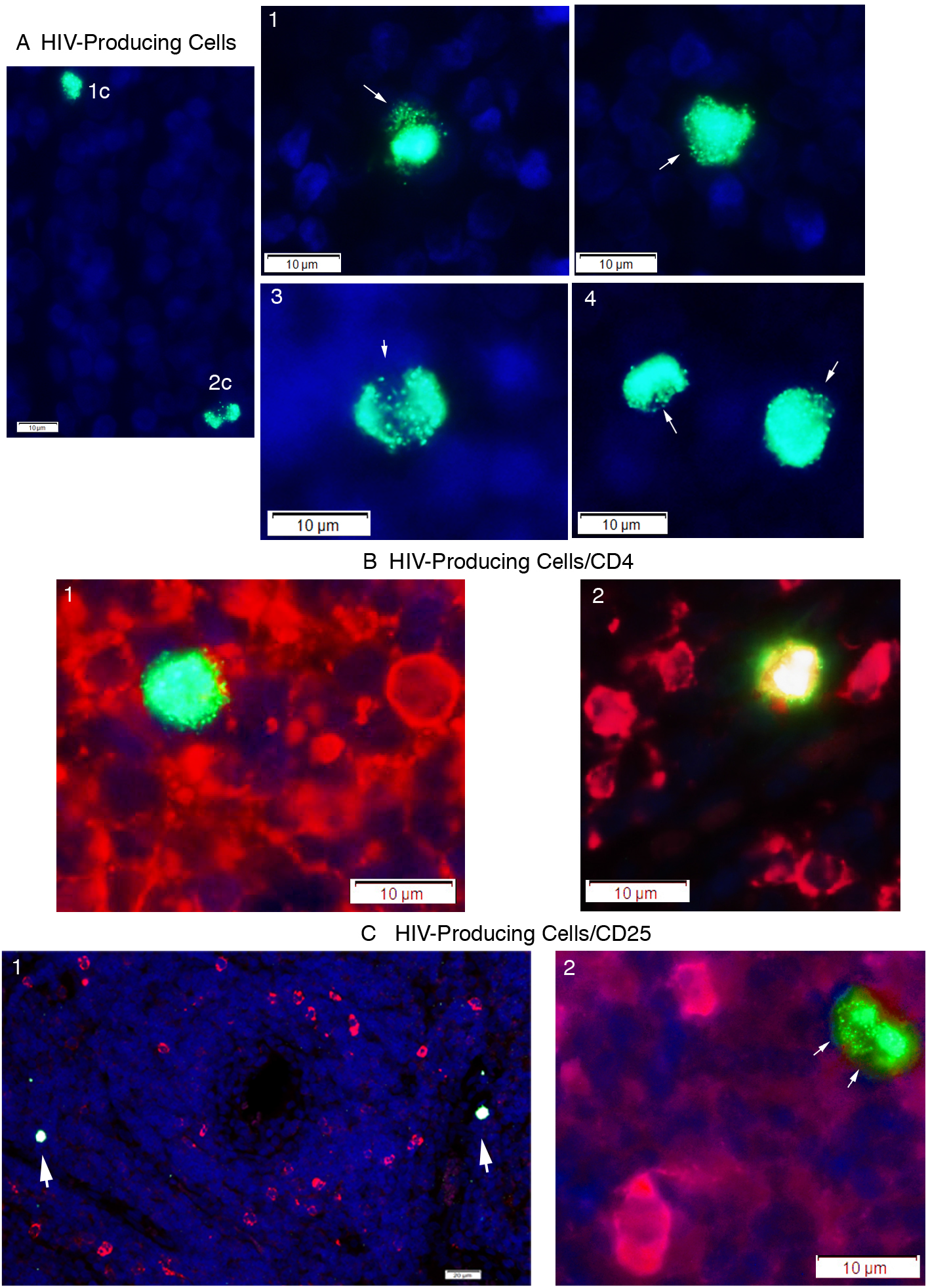
HIV-producing cells in Fiebig 1 are CD4^+^CD25^-^. A. ISH with ELF97 substrate identifies single (1c) or two (2c) productively infected cells together as cells with green virions visible against diffusely stained HIV intra-cellular RNA; DAPI counterstain. A1, 2. Examples of single productively infected cells. A3. Two fused productively infected cells. A4. Two productively infected cells in close spatial proximity. Arrows point to individual and clusters of virions. B. HIV-producing cells are CD4^+^. Virions and intracellular RNA, green; CD4, red. B1.HIV-producing CD4^+^ cell with red/yellow CD4 staining visible at the cell’s margins. B2. Double-labeled red/yellow HIV-producing cell. C. CD25, red; DAPI counterstain. C1. Two CD25^-^ HIV-producing (arrows) cells in a large field with numerous CD25^+^ HIV^-^ cells. C2. Fused CD25^-^ HIV-producing cells (arrows) in the same plane of focus as two CD25^+^ HIV^-^ cells.

We phenotyped the HIV-producing cells by combining ISH with double-label immunofluorescence staining for CD3^+^, CD4^+^ T cells, and CD25^+^ as a marker for activated T cells. In the 8/9 subjects with tissue sections typable by the combined ISH method, 100 percent in 7/8, and, in the remaining 1/8 participant, 98 percent of the HIV RNA^+^ cells were CD3^+^ (Table 1). We found that an average of 96% (range 88-100) of HIV-producing cells were CD4^+^ cells (Figure 1B1, 2 and Table 1) in the 8/9 subjects with typable tissue samples available for the double-label analysis, but only 0.63% HIV-producing cells were CD25^+^ (Table 1 and Figure 1C). We thus attribute virus production in these LN samples in Fiebig 1 to HIV-virus producing CD3^+^/CD4^+^ T cells with a CD25^-^ resting phenotype.

**Figure 2.**
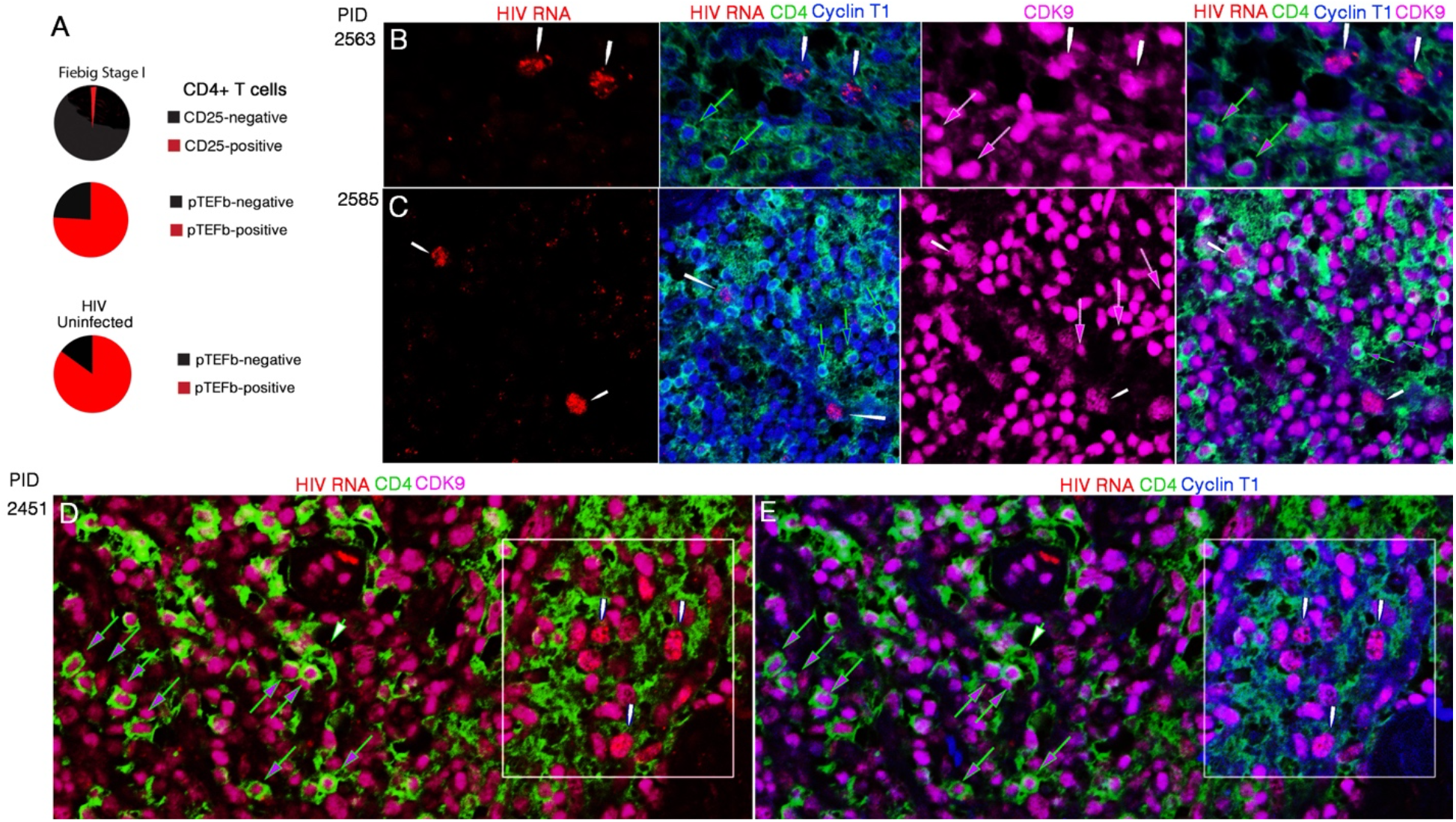
HIV infection of resting CD4^+^pTEFb^+^ populations in Fiebig 1. A. Pie sector diagrams of the CD4^+^CD25^-^ phenotype of 99 percent of CD4^+^ cells, CD4^+^pTEFb^+^ phenotype of 75 percent of CD4^+^T cells in Fiebig 1 and 85% of CD4^+^ T cells in HIV-uninfected LN biopsies. B-E. Images of HIV RNA in CD4^+^pTEFb^+^(Cyclin T1^+^CDK9^+^) T cells in three Fiebig stage 1 participants (PID 2563, 2585 and 2451). White arrows point to location of HIV RNA^+^ cells. B, C. Green outlined blue arrows point to CD4^+^ T cells with Cyclin T1^+^ nuclei. White outlined magenta arrows point to CDK9^+^nuclei. Green outlined blue-magenta arrows point to uninfected HIV RNA-negative CD4^+^ T cells. In the merged image, HIV RNA localizes to CD4^+^ T cells with Cyclin T1^+^ CDK9^+^ nuclei. D. White box encloses HIV RNA^+^ CD4^+^ T cells with CDK9^+^ nuclei. Outside the box, green outlined magenta arrows point to uninfected HIV RNA-negative CD4^+^ T cells with CDK9^+^ nuclei. The green outlined white arrow points to a CD4^+^ T cell with a pTEFb^-^ (CDK9^-^) nucleus. E. Same field as D, except with Cyclin T1-staining in blue. Comparison of the boxed areas in D and E shows that the nuclei of uninfected cells and HIV RNA^+^ CD4^+^ T cells are pTEFb^+^ (Cyclin T1^+^ CDK9^+^).

### Infection of resting T cells in Fiebig 1, target cell availability and pTEFb expression

The principal reason that HIV infects resting T cells is because of their availability: 99 percent of the CD4^+^ T cells available for infection in these Fiebig 1 stage LN samples were CD25^-^ negative (Figure 2A). While that finding would explain why HIV would have to be able to infect resting T cells, it would not explain how the virus could productively infect resting cells that cannot be infected in vitro without stimulation to provide the requisite positive elongation transcriptional factor b (pTEFb)^16-18^. We therefore hypothesized and investigated the possibility that, in vivo, pTEFb is expressed in resting CD4^+^ T cells.

We first confirmed in vitro in PBMC cultures that CDK9 and Cyclin T1 were not detectable by immunofluorescence in cell nuclei in un-stimulated cultures, but both components of pTEFb were detectable after stimulation with CD3CD28 (Supplemental Figure 1). By contrast, in vivo in lymphoid tissues, 85 percent of CD4^+^ T cells in uninfected LN samples from two HIV-uninfected participants in an unrelated study of lymphoid tissue fibrosis, and 75 percent of CD4^+^ T cells in Fiebig stage 1 were pTEFb^+^ (double positive for cyclinT1 and CDK9) (Figure 2A). While the mechanism(s) is/are unknown for this basal state in vivo of pTEFb expression in CD4^+^ T cells, it should provide susceptible target cells to support HIV replication in the lymphoid tissue microenvironment. We documented HIV infection of pTEFb^+^CD4^+^ T cells by combining ISH detection of HIV RNA in CD4^+^ T cells with detection of cyclin T1 and CDK9 together in infected or uninfected cells. In the three Fiebig stage 1 participants we show in Figure 2, HIV RNA co-localizes in CD4^+^ cells with pTEFb^+^ (Cyclin T1^+^CDK9^+^) nuclei in the midst of large numbers of uninfected CD4^+^ cells with pTEFb^+^ (Cyclin T1^+^CDK9^+^) nuclei. In Figure 2B and C, the merged HIV RNA/CD4/Cyclin T1/CDK9 panel shows two HIV RNA^+^ CD4^+^ T cells with Cyclin T1^+^CDK9^+^ nuclei in a field with uninfected CD4^+^ T cells with Cyclin T1^+^CDK9^+^ nuclei. In Figure 2D, we show that nuclei of HIV RNA+ and uninfected CD4^+^ T are CDK9^+^, and, in Figure 2E, that these nuclei are also Cyclin T1^+^.

### Infection of resting T cells in Fiebig 1 and origin of latently infected cells

In the original model of latent infection of CD4^+^T cells, latently infected cells were hypothesized to arise from infected cells that had been activated in an immune response, and, as the cells returned to an immunologically resting state, the proviruses that the cells harbored were silenced^9^. In Fiebig 1, we show here that the only transcriptionally active and productive infections detectable are in resting CD4^+^ T cells, and thus this model cannot account for the origins of a latently infected cell population in the earliest detectable stage of HIV infection. Rather, because we show that HIV replicates in Fiebig 1 in essentially the only target population available of resting CD4^+^ T cells, we hypothesized that the latently infected cell populations in Fiebig 1 LNs are directly established as latent infections in resting CD4^+^ T cells.

To test this hypothesis, we assumed that latently infected cells might be quite rare in Fiebig 1 and therefore devoted the entire ∼10^7^ cells in the LN cell suspensions available to an improved ultrasensitive immunoassay for p24^19^ to maximize the probability of detecting a small reactivatable population of latently infected cells established in Fiebig 1.

We isolated CD4^+^CD25^-^ cells from ∼10^7^ cells from three Fiebig stage 1 participants, one Fiebig stage 4 and one Fiebig stage 5 participant from whom LN suspensions were available. We cultured cells +/- stimulation with CD3/CD28 activator beads and with an integrase inhibitor, and prepared lysates for detection of populations of reactivatable latently infected cells in Fiebig 1. We had previously shown that one reactivated p24^+^ HIV cell in a million uninfected cells produced ∼1.5 pg/ml of p24 in an ultrasensitive immunoassay^15^, and to detect even smaller numbers of reactivated p24^+^ cells in Fiebig 1 we used a combined immunoprecipitation immunoassay with LOD of < 0.005 pg/ml^19^ for a 300-fold increase in sensitivity. We detected p24 production from ongoing replication in unstimulated cells at very low levels close to the LOD that correlated (Spearman’s rank correlation of 0.7) with low levels of detectable HIV RNA^+^ cells (Table 2, Fiebig 1 participants 2391 and 2494, and Fiebig 4, participant 2266). After stimulation, levels of p24 increased five- to more than thirty-fold as evidence of a small p24-reactivatable latently infected CD4^+^CD25^-^ population in Fiebig 1 and Fiebig 4 (Table 2). In Fiebig 1 participant 2540, the high levels of p24 prior to stimulation from ongoing HIV productive infection correlated with increased frequency of HIV RNA^+^ cells, and did not increase after stimulation, presumably because small latently infected populations even after stimulation would be undetectable against the high background of ongoing replication. In this reconstruction, an expanded latently infected population by Fiebig 5 would register after stimulation against high levels of ongoing replication, as was the case (participant 2338, Fiebig 5, Table 2). Thus, while we were limited in availability of samples, as proof-of-principle, HIV demonstrably establishes reactivatable populations of latently infected cells in a target cell population in Fiebig 1 and early-stage infection of CD4^+^CD25^-^ resting T cells.

**Table 2.**
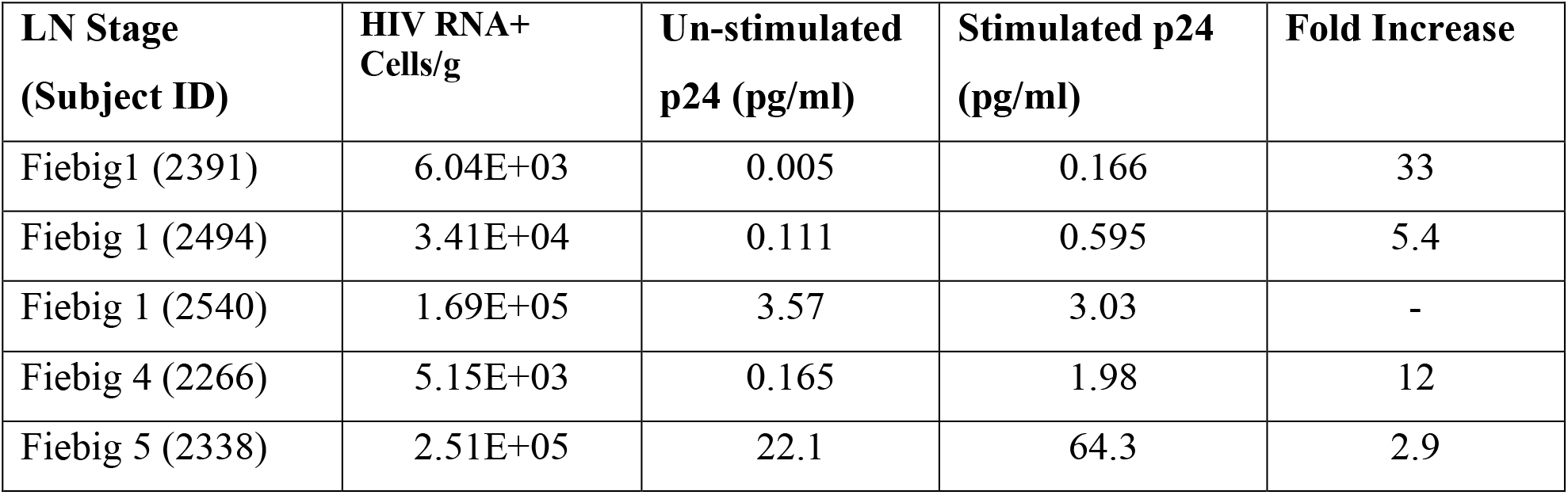
p24 levels in Fiebig 1, 4 and 5 CD4^+^CD25^-^ LN cells +/- stimulation.

| **LN Stage**<br>**(Subject ID)** | **HIV RNA+**<br>**Cells/g** | **Un-stimulated**<br>**p24 (pg/ml)** | **Stimulated p24**<br>**(pg/ml)** | **Fold Increase** |
| --- | --- | --- | --- | --- |
| Fiebig1 (2391) | 6.04E+03 | 0.005 | 0.166 | 33 |
| Fiebig 1 (2494) | 3.41E+04 | 0.111 | 0.595 | 5.4 |
| Fiebig 1 (2540) | 1.69E+05 | 3.57 | 3.03 | - |
| Fiebig 4 (2266) | 5.15E+03 | 0.165 | 1.98 | 12 |
| Fiebig 5 (2338) | 2.51E+05 | 22.1 | 64.3 | 2.9 |

## Discussion

HIV faces the challenge in the earliest stage of infection of replicating in a target cell population almost exclusively limited to resting CD4^+^ T cells. By documenting HIV replication and virus production in these resting CD4^+^ T cell populations in Fiebig 1, we show that HIV like SIV^10-12^ can successfully exploit these initial conditions to generate sufficient virus to sustain and amplify infection as the necessary prelude to establish a persistent infection in the lymphoid tissue reservoir.

We also show that the challenge of infecting resting T cells in vivo is not as daunting as might have been expected from studies of infection in vitro, because a high proportion of CD4^+^ T cells in vivo in lymphoid tissue, operationally defined as CD25^-^ resting T cells, in contrast to resting CD4^+^ T cells in vitro^16^, have readily detectable levels of the critical transcription factor pTEFb. This basal state in vivo in ostensibly resting T cells thus provides HIV with target cells that can immediately support virus production.

Target cell availability in Fiebig1 essentially limited to resting CD4^+^ T cell populations also led us to propose that latently infected resting CD4^+^ T cells most likely originate by directly establishing latent infections in this population, and to show in the reactivation analysis that small latently infected cell populations are in fact established in Fiebig 1 in resting CD4^+^ T cells in lymphoid tissue. We speculate that given the general availability of pTEFb in resting CD4^+^ T cells, the level of Tat and positive feedback control could determine whether a cell is latently or productively infected, as previously described and modeled in vitro^20-22^. Whatever mechanisms decide the fate of the infected resting T cell, the ability to immediately infect resting T cells in Fiebig 1 to generate both latently and productively infected populations enable HIV to persist and propagate in the lymphoid tissue reservoir, and the establishment of a latently infected population at the earliest time HIV infection can be detected and treated also provides one explanation why initiating ART in the earliest possible stage of infection does not prevent rebound when treatment is interrupted^23^. We again conclude that despite the small sizes of the populations of productively and latently infected resting CD4^+^ T cells, strategies beyond initiation of ART in Fiebig 1 will be needed to control or eradicate HIV infection.

## Methods

### Preparation of LN tissue sections

Five-micron sections were cut from 4% paraformaldehyde fixed, paraffin-embedded lymph node samples. Sections of tissues or cell pellets (5 µm) were mounted on Epic Plus Scientific microscope slides (Creative Waste Solutions) and heated at 60°C for 1 hour.

### RNAscope

Every fifth section of the block was dewaxed in xylenes for 10 minutes (twice), placed in 100% ethanol (twice) for 5 minutes before air drying. RNAscope 2.0 Red (ACD) was done as previously published^24^ using the HIV clade AE RNA probe (catalog No. 446551). After mounting, HIV RNA^+^ cells were counted manually.

### RNAscope/Immunohistochemistry

RNAscope was performed using the 2.5 Brown Kit (ACD). After completing amp 6, slides were labeled with Opal 570 for 10 minutes. Slides were microwaved in their appropriate antigen retrieval buffer for 45 seconds at full power before dropping down to 20% power for 10 min. After cooling, samples were blocked in Sniper Blocking solution (Biocare Medical) for 30 minutes before adding the primary antibody, diluted in Divinci Green (Biocare Medical) overnight at 4 degrees. After washing slides in TBST, slides were incubated with opal polymer HRP ms + rab (Akoya biosciences) or HRP-goat (GBI) for 20 minutes. Opal 520 (Akoya Biosciences) was added for 20 minutes before adding DAPI and mounting.

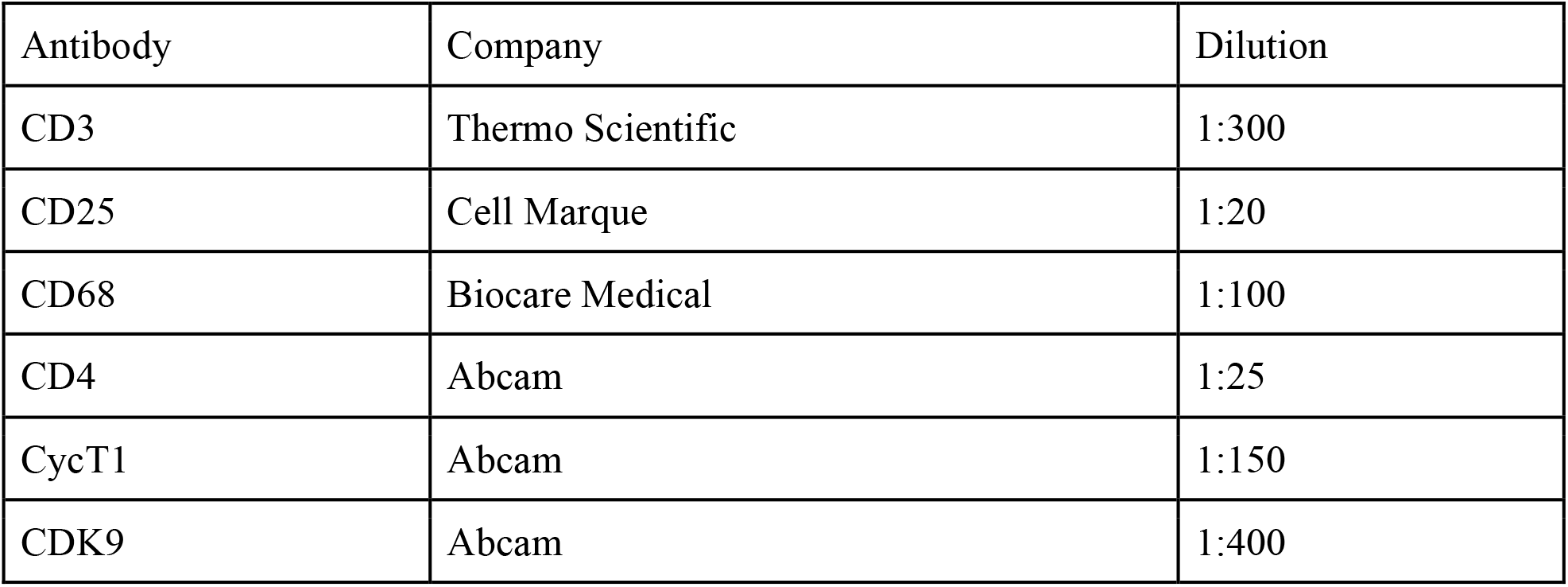

### RNAScope detection of HIV producing cells and phenotypying for CD4 and CD25

HIV-producing cells were detected by incubating tissues at 40°C overnight with AE probe (446551, ACD) and then using the RNAscope 2.5HD detection reagent red kit (322360, ACD) and protocol, except after Amplification 6 step, the tissue sections were washed twice for 2 minutes in wash buffer and then incubated with a 1:10 dilution of ELF97 phosphatase substrate (E6600, ELF™ immunohistochemistry kit, Invitrogen by Thermo Scientific) for 10-20 minutes. In the double label phenotyping analyses, following the ELF97 step, tissues sections were twice washed in TBS for 5 minutes, blocked by background Sniper (BS966L, Biocare Medical) for 45 minutes, before incubating the tissue sections at 4° C overnight with CD4 antibody 1:25 dilution (ab133616, Abcam) or CD25 (4C9) 1:20 dilution (125M-16, Cell Marque). CD4^+^ T cells were subsequently detected by washing the tissue sections twice for 5 minutes in TBS, incubating for 2 hrs at ambient temperature with Alexa Fluor donkey anti-Rabbit 555 1:200 (A31572, Invitrogen), DAPI for 5 minutes and then cover slipping the tissue sections with Aqua-Poly mount (Polysciences). CD25^+^ cells were detected after the TBS wash step by blocking with 3% H_2_O_2_ for 10 minutes, subsequent incubation with Opal Polymer HRP Ms+Rb for 10 minutes, washing twice for 5 minutes in TBS, Opal 570 reagent 1:100 (NEL810001KT, Akoya Biosciences) for 10 minutes, washing, DAPI staining and mounting as described for CD4. Slides were stored at 4 ° C until analyzed as follows: The entire section was scanned under WU illumination at 40X to first detect and capture images of HIV-producing cells followed by images of CD4^+^ or CD25^+^ cells at the same location using the red filter. Images of the ELF97-HIV-producing cells and CD4 or CD25 were merged in Photoshop and single positive HIV-producing and double positive CD4^+^ or CD25^+^ were enumerated manually.

### Detection of HIV RNA^+^ pTEFb^+^ CD4^+^ T cells

Immunofluorescence staining for HIV RNA, CycT1, CDK9, and CD4 proteins was performed simultaneously on 6um thick LN sections by a modification of Zhang et al.^11^. Slides were dewaxed, treated with 1 ug/ml proteinase K, and hybridized to HIV anti-sense RNA probe labelled with digoxigenin. After TSA and Opal 620 to detect HIV RNA, multiplex Fluorescent IHC was performed using an Akoya Opal Multiplex IHC kit. Sections were heated for a minimum time between steps to remove previous antibodies and unmask the next antigen. CD4 (Opal 520) was followed by CycT1 (Opal 690) and then CDK9 (Opal 405). Each antigen was unmasked using its optimal buffer. 3-5 sections were tested for each tissue and patient, and images were taken using a Zeiss LSM800 Confocal/Super-resolution Microscope at 20X. Each CD4 cell in a field was numbered and scored for the presence of nuclear CycT1 and CDK9 using single color channel overlays.

### Detection of reactivatable latently infected CD4^+^CD25^-^ T cells by ultrasensitive p24 immunoassay

Isolation of CD4^+^CD25^-^ cells from frozen suspensions was performed using negative selection with a Miltenyi CD4 human T cell isolation kit to which biotinylated CD25 was added in excess.^25^ Cells were cultured at 10^6^ cells/ml for 6 days in RPMI containing 100nM raltegravir. Half of the cells were stimulated with Human T Cell activator CD3/CD28 beads at a bead/cell ratio of 1:1. After 3-5 days, both cell cultures were lysed in PBS containing 0.5% Triton X 100 and 3% BSA, and the stimulated and unstimulated lysates tested for HIV p24 by combined immunoprecipitation and digital ELISA^19^.

## Acknowledgments

The authors thank Tim Leonard and Colleen O’Neill for assistance with preparation of the figures and manuscript. The study team is grateful to the individuals who volunteered to participate in this study and the staff at SEARCH/IHRI, the Thai Red Cross AIDS Research Centre and the Department of Retrovirology, U.S. Army Medical Component, Armed Forces Research Institute of Medical Sciences (AFRIMS), and the RV254/SEARCH 010 Research Team. We are grateful to the Thai Government Pharmaceutical Organization (GPO), ViiV Healthcare, Gilead Sciences and Merck & Co., Inc., Rahway, New Jersey, USA for providing the antiretroviral medications for the RV254/SEARCH 010 study.

## Author Contributions

ATH, TWS, PJS and SW designed the studies, developed and validated the methodology, and analyzed the results. SW, JA, and LD designed and performed the image analyses. SW, PZ and BJH contributed to experimental design and access to new assays/tools and GW performed the reactivation studies. CR provided statistical analysis. EK, CS, NT, DC, NC, PP, SP, DH, SV, MR, NP and JA managed participant recruitment, enrollment, follow-up and overall conduct of the RV254 cohort. SC, SB, RT, SM, MdS, ST, AS managed specimen collection, processing and laboratory assays involved in Fiebig staging.

ATH wrote the paper with contributions from all co-authors.

## Competing Interest Declaration

GW, PZ and BJH are employed by Merck Sharp & Dohme LLC, a subsidiary of Merck & Co., Inc., Rahway, New Jersey, USA.

### Disclaimer

The views expressed are those of the authors. The content of this publication does not necessarily reflect the views or policies of the Department of Health and Human Services, U.S. Army or Department of Defense, nor the Henry M. Jackson Foundation for the Advancement of Military Medicine, Inc., nor does mention of trade names, commercial products, or organizations imply endorsement by the U.S. Government including the U.S. National Institutes of Health. The investigators have adhered to the policies for protection of human subjects as prescribed in AR-70-25.

## Supplementary Information

**Supplemental Table 1.**

**A. Participant stage, demographics, CD4 counts, pVL and clade**
| **PID** | **Fiebig Stage** | **Estimated Time (Days) from HIV Exposure** | **Age** | **CD4** | **pVL** | **Clade** |
| --- | --- | --- | --- | --- | --- | --- |
| 2265 | 1 | 13 | 22 | 354 | 8,970 | AE |
| 2391 | 1 | 18 | 26 | 575 | 11,969 | AE |
| 2397 | 1 | 12 | 23 | 654 | 31,389 | AE |
| 2246 | 1 | 4 | 21 | 371 | 18,311 | AE |
| 2540 | 1 | 26 | 28 | 572 | 37,142 | AE |
| 2451 | 1 | 17 | 27 | 699 | 5,360 | AE |
| 2563 | 1 | 12 | 27 | 405 | 586 | AE |
| 2585 | 1 | 11 | 36 | 493 | 35,255 | AE/B Recombinant |
| 2494 | 1 | 16 | 28 | 761 | 8,561 | AE |
| 2266 | 4 | 22 | 34 | 519 | 100,793 | AE |
| 2338 | 5 | 44 | 39 | 625 | 277,506 | AE |

**B. Fiebig staging systems used to classify study subjects with acute HIV infection**
| **Staging System** | **Stage** | **Serological Characteristics** |
| --- | --- | --- |
| Fiebig | 1 | NAT+ /p24 antigen- /HIV IgM-, IgG-, WB- |
|  | 2 | NAT+ /p24 antigen+ /HIV IgM-, IgG-, WB- |
|  | 3 | NAT+ /p24 antigen+ /HIV IgM+, IgG-, WB- |
|  | 4 | NAT+ /p24 antigen+/- /HIV IgM+, IgG -, WB indeterminate |
|  | 5 | NAT+ /p24 antigen+/- /HIV IgM+, IgG +/-, WB+ without p31band |

Abbreviations: NAT, nucleic acid testing; WB, Western blot

**Supplemental Figure 1.**
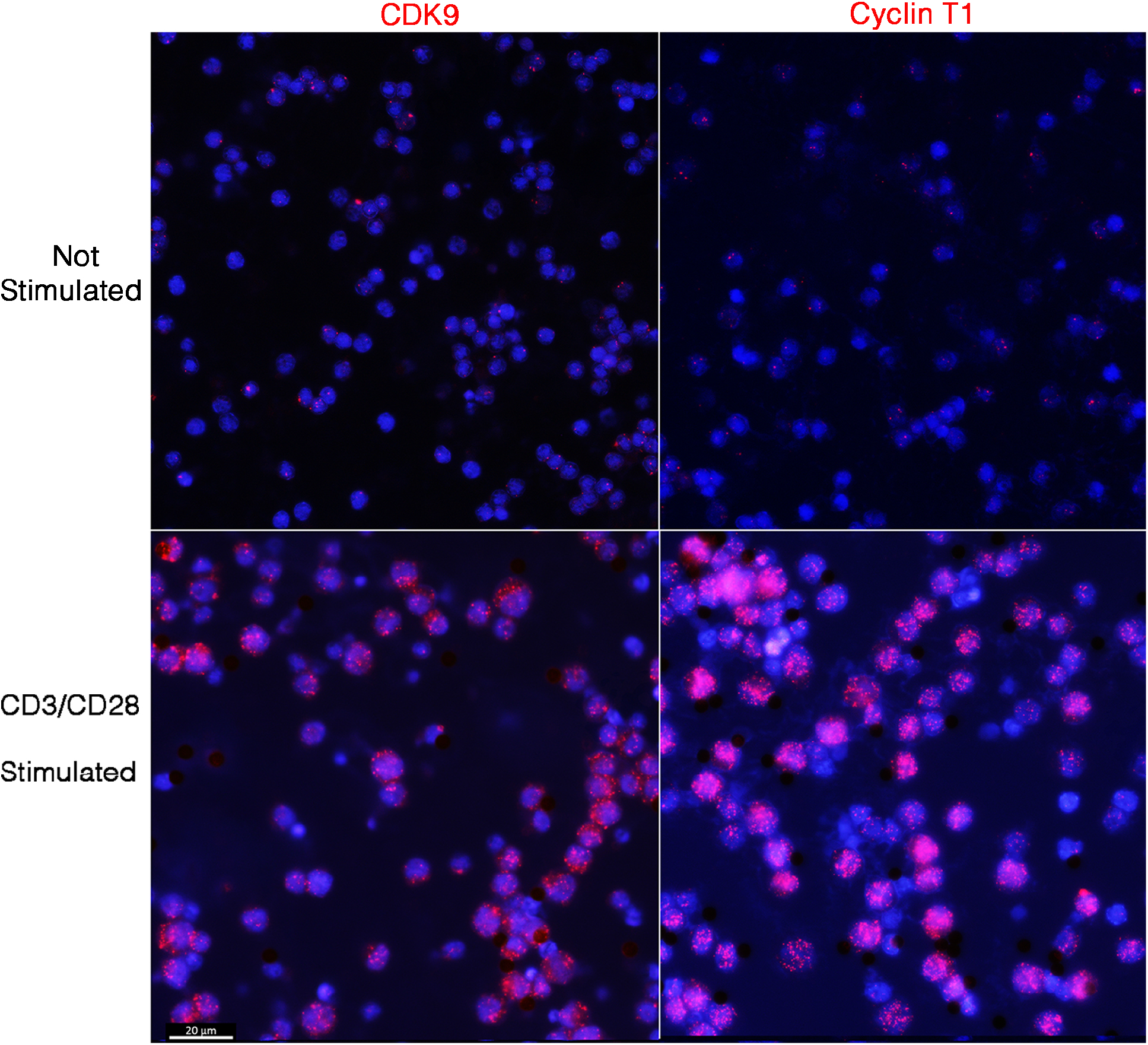
In vitro expression of CDK9 and Cyclin T after Stimulation. PBMC cultures, either not stimulated or stimulated with CD3CD28 activator beads, were stained with antibodies to CDK9 (left panels) or Cyclin T1. Nuclei counterstained blue with DAPI. Without stimulation, CDK9 or Cyclin T1 are not detectable in the nuclei. Cell stimulation induces expression of both CDK9 and Cyclin T1 in a speckled pattern in the nuclei of stimulated cells. Scale bar for all four panels, 20 µm.

